# Label-free three-dimensional observations and quantitative characterisation of on-chip vasculogenesis using optical diffraction tomography

**DOI:** 10.1101/2020.01.01.892620

**Authors:** Chungha Lee, Seunggyu Kim, Herve Hugonnet, Moosung Lee, Weisun Park, Jessie S. Jeon, YongKeun Park

## Abstract

Label-free, three-dimensional (3D) quantitative observations of on-chip vasculogenesis were achieved using optical diffraction tomography. Exploiting 3D refractive index maps as an intrinsic imaging contrast, the vascular structures, multicellular activities, and subcellular organelles of endothelial cells were imaged and analysed throughout vasculogenesis to characterise mature vascular networks without exogenous labelling.

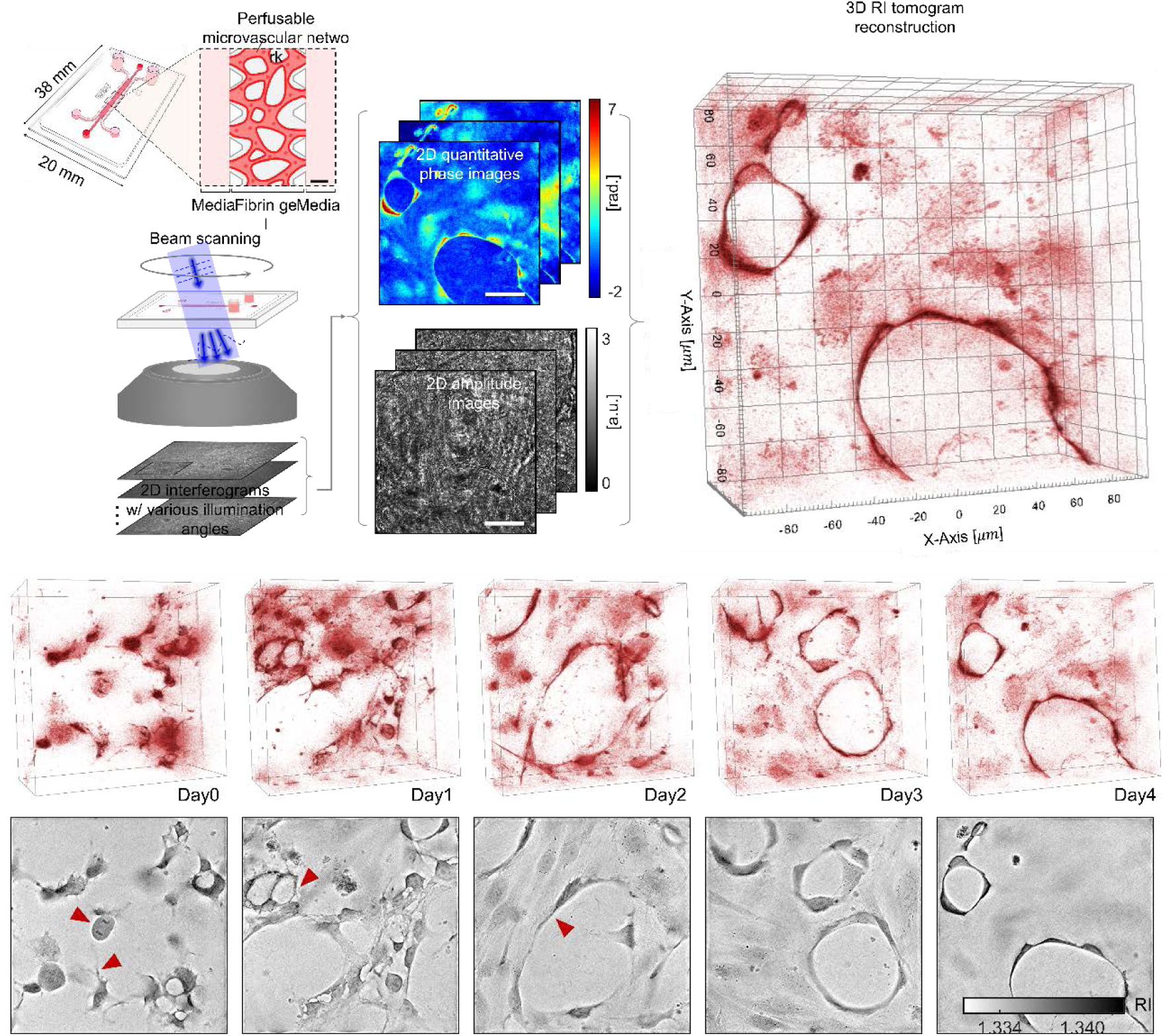

---

Significant breakthroughs in on-chip miniaturisation have recently been achieved with diverse potential applications, including three-dimensional (3D) multicellular micrometre-scale bioassays and drug discovery. These advanced *in vitro* platforms have overcome several shortcomings of conventional two-dimensional (2D) static culture plates, making it possible to reconstitute *in vivo* microenvironments and functions, as displayed by investigations of the vasculature, gut, eye, and cancer metastasis^1-4^.

Despite this significant progress in on-chip technology, the observations and assessments of cells and their architectures in on-chip devices mostly require the use of labelling agents such as fluorescence proteins or organic dyes. Various imaging techniques have been utilised for these purposes, including epi-fluorescence microscopy^5, 6^, phase-contrast microscopy^7, 8^, and confocal fluorescence microscopy (CFM)^7,9^. In particular, CFM is a well-established imaging technique with advantages owing to its 3D imaging capability and high molecular specificity^10^. However, the requirement of fluorescent labelling procedures introduces several drawbacks such as phototoxicity and photobleaching^11^ that ultimately limit the long-term imaging of 3D multicellular dynamics inside a microfluidic chip. Therefore, overcoming this limitation could help to propel lab-on-chip approaches in more diverse applications of biotechnology and medical science to fully exploit the advantages of 3D multicellular culture and targeted assays.

To address these issues, we here report the successful label-free 3D observations and quantification of 3D multicellular culture on chip devices. Specifically, we exploited the refractive index (RI) distributions of cells, as an intrinsic optical property of materials, to achieve the label-free and quantitative bioimaging of 3D multicellular structures in on-chip vasculogenesis. The 3D-cultured endothelial cells were systematically investigated at subcellular resolutions using optical diffraction tomography (ODT), a 3D quantitative phase-imaging (QPI) technique, which reconstructs the 3D RI distributions of a sample from multiple 2D interferograms measured at various illumination angles^12^. The 3D tomographic imaging capability of ODT has been demonstrated for various types of biological samples to date, including red blood cells^13^, neuron cells^14^, white blood cells^15^, cancer cells^16, 17^, and bacteria^18^. Recently, QPI was applied to microfluidic technologies with synergistic experimental applications for imaging floating cells^19-21^, and for investigating microfluidic mixing^22, 23^ and angiogenic sprouting^24^. However, on-chip multicellular behaviours that appear in a truly 3D manner have yet to be explored using 3D QPI techniques.

The experimental timeline (Fig. 1A, right) consisted of three steps: 1) chip preparation, 2) 3D observation of on-chip vasculogenesis for 4 days from the day of cell seeding, and 3) characterisation of the day-4 mature microvascular networks via quantification of its structure and permeability. First, microvascular networks inside a polydimethylsiloxane (PDMS) microfluidic chip were constructed over 4 days in a 3D fibrin gel, based on a previously optimized protocol^25^ (ESI†). In brief, the microfluidic chips were prepared by bonding a 1.5-H-thick cover glass to the micropattern-engraved surface of the thin PDMS chip, and two perforated PDMS blocks were then bonded to the unbonded surface of the PDMS to supply abundant cell culture media and prevent drying (Fig. 1A, left). To provide a sufficient working distance for the high numerical aperture (NA) objective lens, the thickness of the PDMS chips was set to 1 mm, which is relatively thinner than applied in typical chip fabrication.

**Fig. 1.**
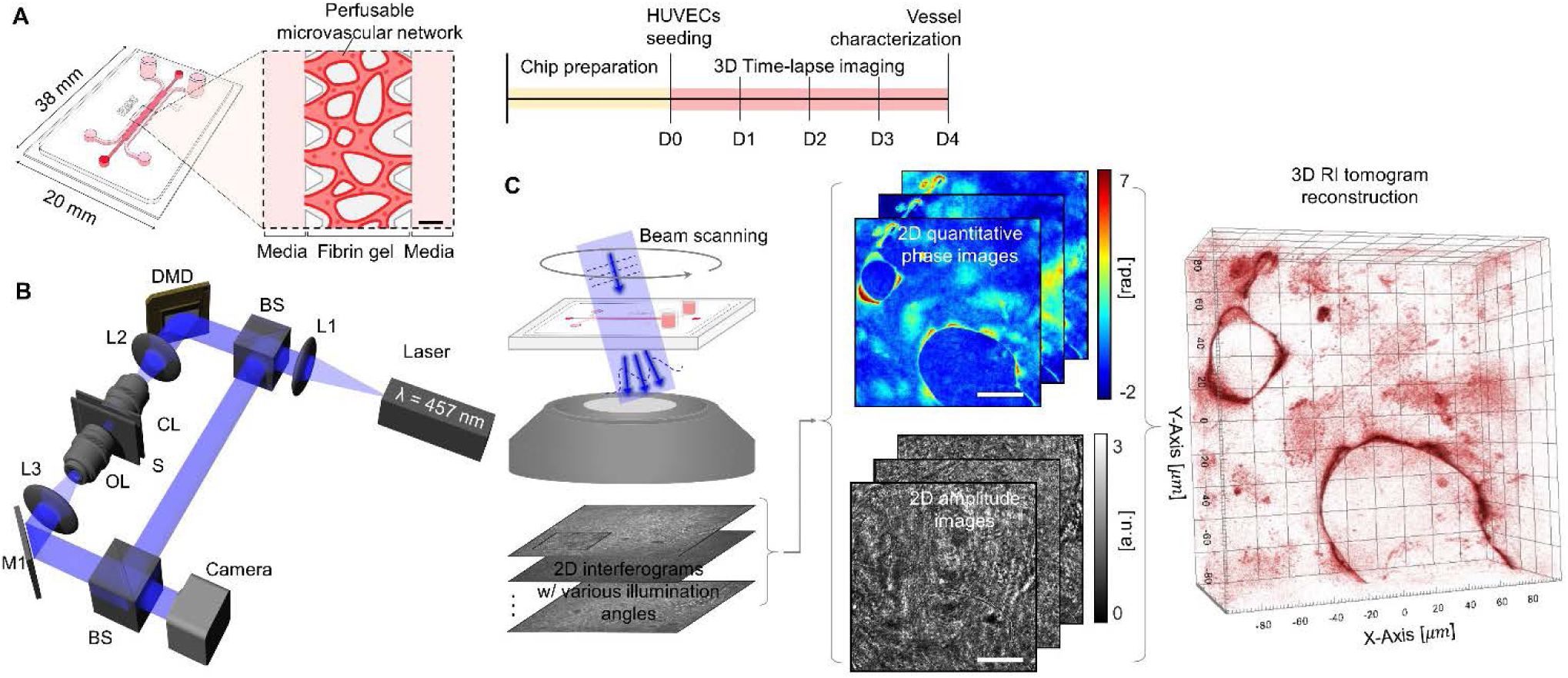
Schematic diagrams of the experimental procedures. (A) Microfluidic chip fabrication (left) and experimental timeline (right). Scale bar = 300 µm. (B) Experimental setup of ODT: L1-L3, lenses (*f*_*1*_ = 400 mm; *f*_*2*_ = 175 mm; *f*_*3*_ = 500 mm); BS: beam splitter; DMD: digital micromirror device; CL: condenser lens; S: sample; OL: objective lens. (C) Flowchart of ODT: 1) measurement of 2D multiple interferograms; 2) retrieval of 2D scattered fields using the 2D interferograms; 3) reconstruction of the 3D RI tomogram of the sample. Scale bar = 50 µm.

To confirm vasculogenesis, human umbilical vein endothelial cells (HUVECs) transfected with red fluorescence protein (Angio-Proteomie) were mixed with a fibrin gel (6 mg/mL) and the mixture was immediately injected to the chip. The HUVECs in the 3D fibrin gel were then cultured at 37°C in a 5% CO_2_ incubator in EGM2MV (Lonza) medium supplemented with exogenous vascular endothelial growth factor-A (50 ng/mL, Peprotech). The medium was changed daily.

A custom ODT system was constructed for 3D observation of the HUVECs forming microvascular networks (Fig. 1B). The ODT setup was based on a Mach-Zehnder interferometer equipped with a digital micromirror device (DMD). A blue continuous-wave diode-pumped solid-state laser (wavelength = 457 nm, Cobolt Twist, Cobolt, Sweden) served as a coherent light source. The DMD enables high-speed and high-stability beam scanning for sample illumination^26, 27^. The use of a long-working-distance (0.80–1.80 mm) condenser objective (LUCPLFLN40X, NA = 0.6, Olympus Inc.) allowed for beam focusing within a sample plane penetrating the 1-mm PDMS. The scattering light from a sample was collected by an objective lens (UPlanSAPO20X, NA = 0.75, Olympus Inc.) and then projected to an image plane, where both the sample and reference beams interfere to form a spatially modulated interferogram. Interferograms of a sample were recorded using a CMOS camera (LT425M-WOCG, Lumenera Inc.) with the frame rate set to 60 Hz.

Owing to its label-free imaging capability, ODT enabled the long-term (5 days) 3D imaging of the same device (Fig. 2). 3D multicellular activities were observed while the multiple HUVECs in the fibrin gel formed lumenised vasculatures, including cell division, apoptosis, filopodia extension, vacuoles formation and fusion, luminal expansion, and final microvessel maturation (Fig. 2A, filled arrows, and S1, ESI†)^28, 29^.

**Fig. 2.**
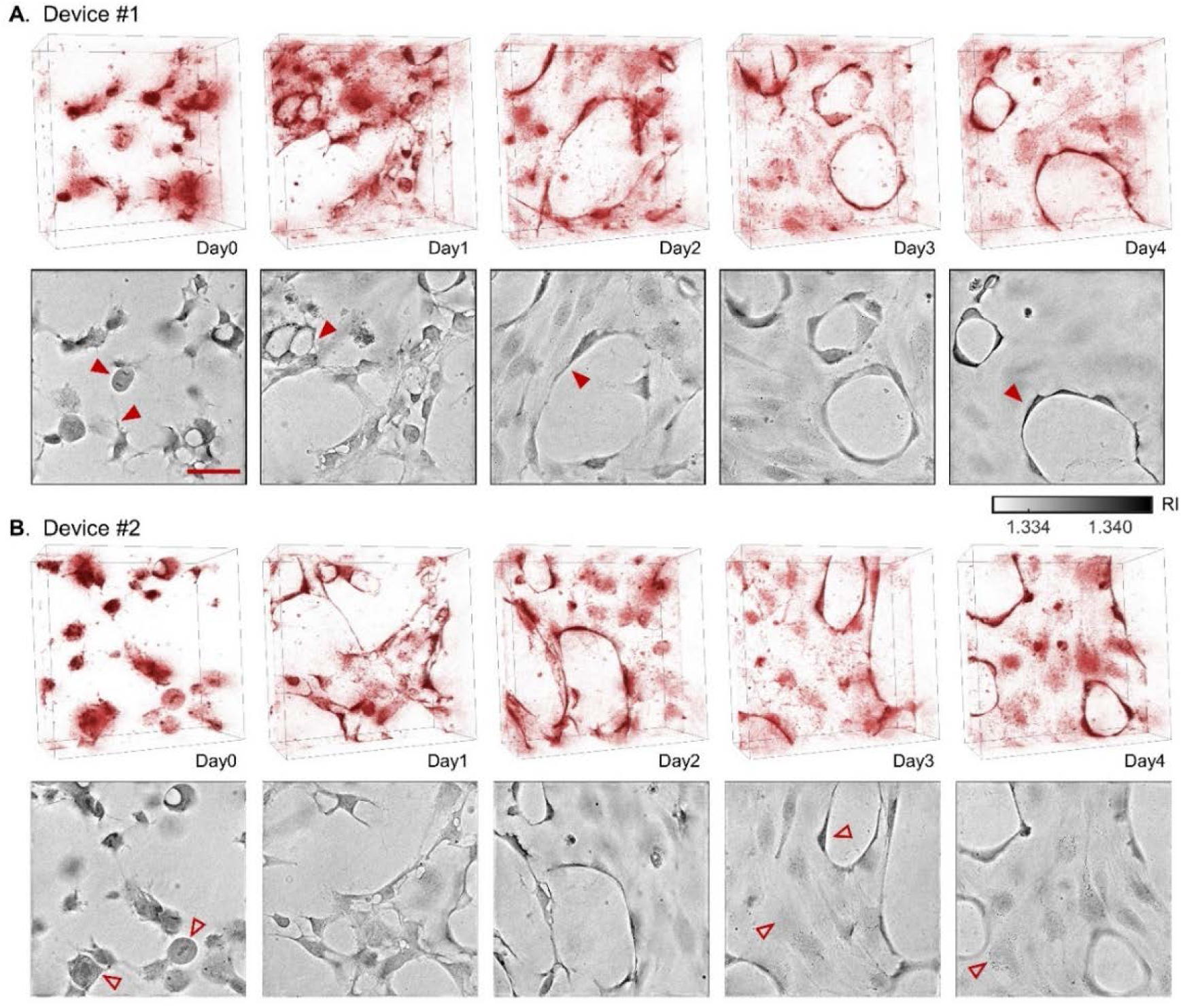
Three-dimensional observation of on-chip vasculogenesis using ODT. (A-B) Three-dimensional rendering (top) and a representative XY-slice image (bottom) of 3D RI tomograms of 4-day on-chip vasculogenesis for the two selected devices, A and B, respectively, in which various cellular activities (filled arrows) and subcellular organelles (open arrows) were observed during vasculogenesis. Scale bar = 50 µm.

The 3D RI tomogram of the sample was then reconstructed from the multiple 2D holograms of a sample measured with various illumination angles (Fig. 1C). A total of 109 2D holograms were recorded for each sample, from which 2D scattered fields of phase and amplitude information were retrieved using a field retrieval algorithm^30, 31^. The 3D RI tomogram was then reconstructed from these scattered fields based on the Fourier diffraction theorem^32^. Owing to the limited NAs of both the condenser and objective lens, side-scattering signals were not collected, resulting in degradation of the axial resolution in ODT. To address this issue, known as the missing-cone problem, an iterative regularization algorithm was used^33^. The RI value of the cell culture medium was set to 1.334, which was experimentally measured using a refractometer at room temperature (Atago™ 2350, Thermo Fisher Scientific Inc.).

The theoretical resolution of our imaging system is 0.19 and 0.95 μm in the lateral and axial directions, respectively^34^. Image reconstructions, analysis, and visualisation were performed using MatLab (MathWorks Inc., Natick, MA, USA), ImageJ (NIH, USA), and Tomostudio (Tomocube Inc., Republic of Korea). The details on the principles of ODT, and the 3D reconstruction MatLab codes can be found elsewhere^13, 35^.

The high-resolution imaging capability (190 nm) of the system enabled identification of the subcellular organelles of individual cells in the reconstructed 3D RI tomograms, such as the nucleoli, chromosomes, microtubules, actin filaments, Golgi apparatus, and liposomes (Fig. 2B, open arrows, and S1, ESI†). Of note, ODT can obtain images of an object with rapidly changing morphologies since the measured field scattered from the object truly mirrors its spatial distribution. Therefore, the vasculogenesis of HUVECs, which involves dramatic changes in cell shapes and morphologies, can be further quantitatively investigated using ODT. This approach can be readily extended to reveal the underlying mechanism in cell-matrix remodelling during vasculogenesis. Therefore, after 4-day on-chip vasculature, the mature microvascular networks were further characterized using ODT. For cross-validation purposes, we first compared the results obtained with ODT with those obtained using the well-established CFM approach (Zeiss LSM880, an objective lens of NA = 0.8 with z-step of 2.1 μm). For a fair comparison, we acquired images in the same field-of-view (641 μm × 641 μm) of the same microfluidic device using both CFM and ODT after 15 min of 4% paraformaldehyde fixation at room temperature. Multiple 3D images that covered the same region of interest (ROI), i.e., 5 × 5-tiled images for ODT and 2 × 2-tiled images for CFM, were acquired, stitched, cropped, and registered using the built-in *imregister* function in MatLab. With respect to revealing the overall 3D structure of the mature microvascular networks, ODT was found to be comparable to CFM (Fig. 3A and 3B). The endothelium of the microvascular networks could be clearly distinguished in the Z-projection images for both CFM and ODT, along with endothelial cells forming aligned vessels, suggesting that both techniques are capable of investigating the overall architecture of on-chip vasculatures.

**Fig. 3.**
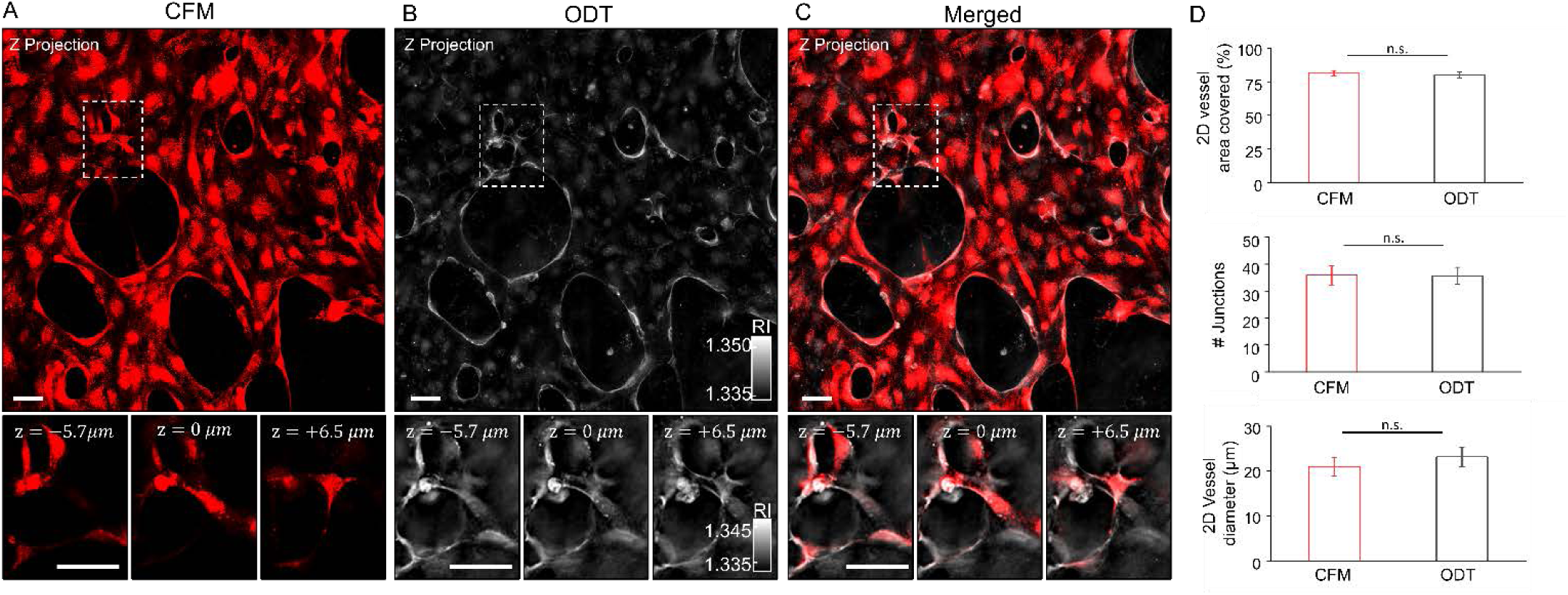
Characterisation of mature microvascular networks using both CFM and ODT for comparison. (A-B) Z-projection images of the same field-of-view (641 μm × 641 μm) acquired by CFM (A) and ODT (B), in which the dashed boxes of the same ROI are presented in the zoomed-in XY images at z= −5.3 μm (left), 0 μm (middle), and +6.5 μm (right) below the Z-projection images. (C) Merged image of (A) and (B). (D) Quantitative characterisation of the microvascular architectures with respect to the vessel area (top), junction numbers (middle), and vessel diameter (bottom). Scale bar = 50 µm. Error bars = SEM.

Furthermore, multiple XY images at three Z-axis positions (dashed boxes in Fig. 3A and 3B) reflected the 3D nature of the vessel geometry, which could be barely detected in the Z-projection images. These results suggest that ODT is a more powerful imaging technique for not only the long-term imaging of the on-chip vasculature process but also for characterisation of 3D vessel geometry. Despite the apparent similarity in the overall microvessel architecture, fundamental differences between ODT and CFM also influence the capabilities of these methods. Intuitively, a colour bar is not available for analysis of the CFM image. Moreover, ODT can further provide the 3D RI distribution of the microvascular network, which has been well-recognized as a key biophysical parameter in microfluidic technologies^20^.

A merged image of the Z-projection images obtained by CFM and ODT showed relatively high correlation (Fig. 3C). The comparable information on the overall vascular architecture provided by ODT further enabled obtaining quantitative structural information of the mature vascular networks, including the vessel area, junction numbers, and vessel diameter (Fig. 3D, and ESI†). We compared a total of nine images of three different ROIs inside three different microfluidic devices, and no significant differences between the two methods were found.

To further demonstrate the quantitative imaging capability of ODT, we quantified the microvascular permeability of the endothelium of the on-chip vasculature. The microvascular permeability coefficient *P* is an important physiological index for evaluating the endothelial barrier function^36^. To date, fluorescence imaging techniques have been utilised in microvascular permeability assays in which fluorescence emission from macromolecules such as fluorescein isothiocyanate-dextran is detected during diffusion through an endothelium of a microvessel^4,37^. As an alternative label-free approach, by exploiting the distinct RI values of dextran molecules, we successfully observed and quantified the diffusion of unlabelled dextran macromolecules at a frame rate of 1 Hz. For quantification of *P*, we measured the optical phase delay *Δϕ* as an integration of RI values over an optical path length. Pulse-like injection of the dextran solution will result in intravascular perfusion of the dextran molecules, followed by its diffusion through the endothelium, and then *Δϕ* can be measured (Fig. 4A). For normalisation, the reported *Δϕ* was obtained after subtraction of the box-averaged (10 pixels squared) value of the hydrogel and intravascular regions, which reduces the effect of random phase fluctuation (Fig. 4D, inset at the lower left) and cancels out a global phase fluctuation in both regions as an arbitrary constant value. The day-4 mature microvascular networks were measured for 10 min, followed by injection of 50 μL dextran solution [110-kDa dextran macromolecules (Dextran T110, Pharmacosmos) dissolved in phosphate-buffered saline at 200 mg/mL] into the microfluidic channel. As a result, the phase images at five selected time points mirrored the actual diffusion corresponding to the schematic in Fig. 4A (Fig. 4B, movie, ESI†). Intuitively, the transendothelial diffusion of the dextran macromolecules is responsible for the so-called RI matching effect, by which the increased optical phase delay (OPD) in a gel region gradually reduces the phase difference between the gel and intravascular region (Fig. 4B, the 4^th^ and 5^th^ images from the left). The kymograph of the dashed line aa’ in Fig. 4B effectively visualises the RI-matching effect at the three different vessel–gel interfaces, at which the phase difference of the two regions gradually decreases, thereby smoothing the boundaries during transendothelial diffusion (Fig. 4C).

**Fig. 4.**
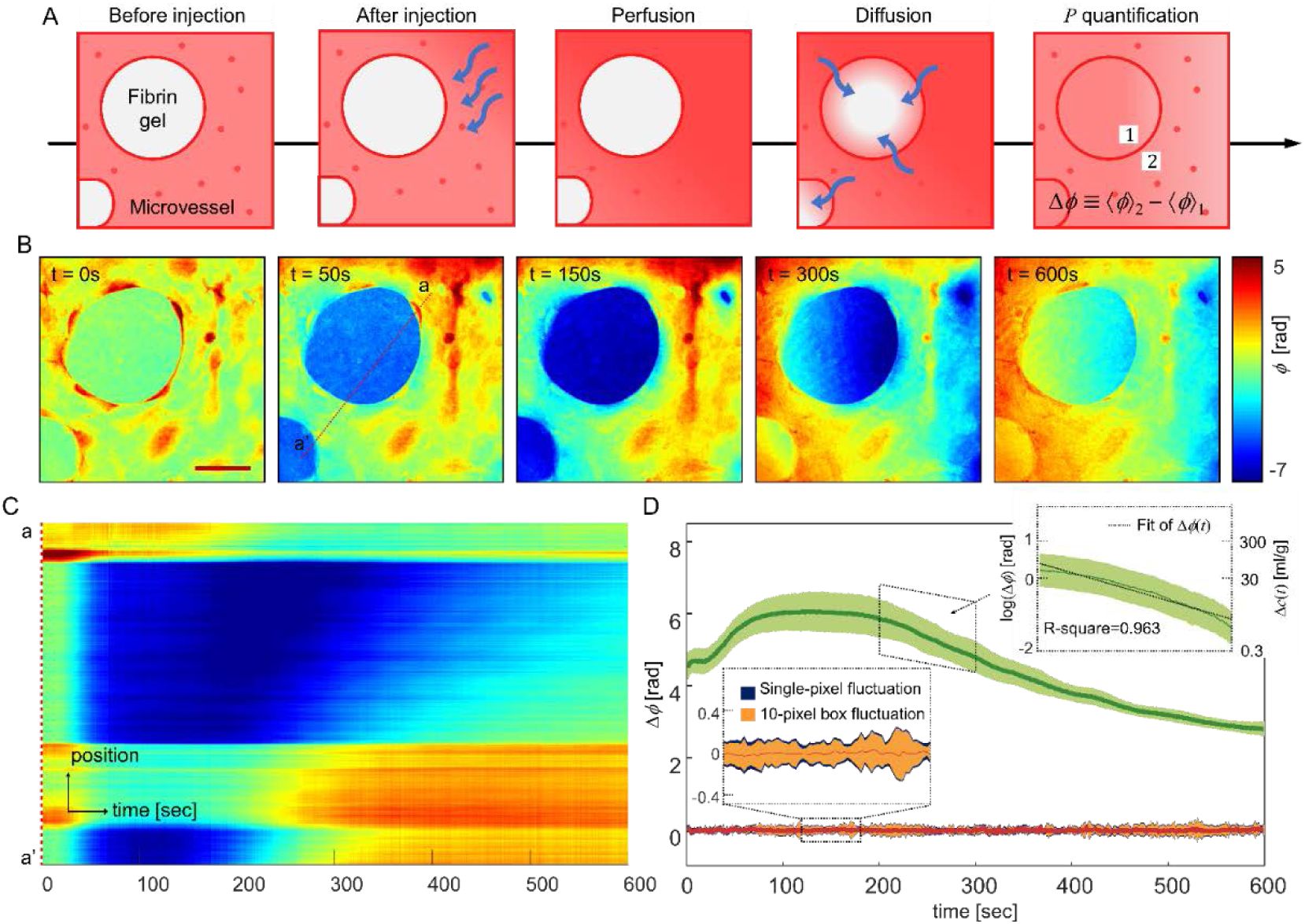
Phase-based microvascular permeability assay using ODT. (A) Schematic diagram of the microvascular permeability assay with a pulse-like injection of 110-kDa dextran (1-2) that is monitored through perfusion and transendothelial diffusion (3-4), allowing for *P* quantification with Δ*ϕ*(*t*) (5) defined as the subtraction of the 10-pixel box average of an intravascular- and gel-region. (B) The measured diffusion data corresponding to the schematic (A), in which the entire dynamic process inside the chip was recorded every second for 600 s. Scale bar = 50 µm. (C) Kymograph of the dashed line aa’ in (B), which penetrates three different vessel–gel interfaces. (D) Validation of the phase stability of the ODT system and *P* quantification. The stability data from the no-sample measurement (orange) and the actual diffusion data (green) presented as the average (line) and standard deviation (shading bar), respectively. The inset at the lower left shows a reduced random phase fluctuation with 10-pixel box representation of Δ*ϕ*(*t*). The inset at the upper right shows *P* quantification from the fit function of Δ*ϕ*_*d*_(*t*).

Based on the measured images, *P* was further quantified with the manually calculated *Δϕ(t)* from 15 different ROIs (Fig. 4D). Before *P* quantification, we first validated the phase stability of ODT. The stability data acquired from the no-sample measurement confirmed the random phase fluctuation of *Δϕ*(*t*) = −0.01 ± 0.35 radian, which was clearly distinct from the actual diffusion data (Fig. 4D, orange and green, respectively, with the average and standard deviation presented as a line and shading bar). For estimation of *P* with phase information, the widely used formula of *P* was slightly modified ^38,39^ (ESI†). Intuitively, we exploited the linear relation of the OPD induced by the diffusing dextran macromolecules, *Δϕ*_*d*_*(t)*, and its concentration difference, *Δc(t)*^40^. From the fit function of *Δϕ*_*d*_*(t)*, the estimated value of *P* was 8.874 × 10^−6^ cm/s with R^2^ of 0.963. Considering the variability of biological samples, we conclude that the order of the measured value is consistent with the reported permeability values of 70-kDa dextran across the 3D microvasculature (2.180 ± 0.2277 × 10^−5^ cm/s)^41^ and 2D endothelial monolayer (3.9 ± 0.7 × 10^−6^ cm/s)^42^ determined on *in vitro* platforms.

Overall, these results demonstrate the ability of an ODT approach to achieve 3D label-free observation and characterisation of on-chip vasculogenesis. The 3D RI tomograms provided volumetric information about on-chip vasculogenesis, which was compatible with that obtained using conventional fluorescence confocal microscopy, but without requiring any exogenous labelling agents. The label-free imaging capability was highlighted by measuring on-chip vasculogenesis for 5 days without needing to change the sample. Moreover, by exploiting the high spatial and temporal resolution of ODT, the subcellular imaging of complex multicellular behaviours and quantitative diffusion analysis were realized.

Despite these clear advantages of label-free and quantitative imaging, one of the main limitations of ODT is its lack of molecular specificity. Therefore, although the RI tomograms provide valuable information about the structures and cells during vasculogenesis, identifying the contributions of specific molecules or proteins in these processes still requires labelling approaches. Thus, a combined approach of RI and 3D fluorescence signals would allow access to both structural and molecular information^43-45^. Another limitation is that on-chip vasculogenesis was measured from the 3D RI tomograms, whereas the quantification of *P* was performed from the 2D-projected RI images. This was mainly due to the limited speed of the image sensor, because the speed of ODT is not limited by the illumination control but rather by the image acquisition. Therefore, faster 3D quantifications of *P* could be realized by employing a high-speed image sensor^46^.

Although we demonstrated the 3D label-free quantitative imaging approach for on-chip vasculogenesis, this system can be readily expanded to various applications, including 3D cell culture, organ-on-a-chip, organoid, and drug screening and discovery. The proposed method would also be applicable for further investigating the dynamic behaviours of multiple cells when interacting with their 3D microenvironment *in vitro*. Given the rapid advances in organ-on-a-chip technology, we believe that ODT will emerge as a powerful imaging technique for exploring microfluidic 3D cell culture models in a label-free and quantitative manner.

## Author contributions

Y.K.P. and J.S.J. initiated the work and supervised the project. C.L. and S.K. performed the experiments. H.H. and H.L. developed the methods. All authors wrote the manuscript.

## Acknowledgements

This work was supported by KAIST, BK21+ program, Tomocube, the K-valley RED&B (Grant No. N11190070) of the KAIST, and National Research Foundation of Korea (2017R1D1A1B03030428, 2017M3C1A3013923, 2015R1A3A2066550, 2018K000396).

