## Supplementary Information for "Label-free three-dimensional observations and quantitative characterisation of on-chip vasculogenesis using optical diffraction tomography"

**Microfluidic chip fabrication and vascular networks formation**

We created 3D vascular networks in a hydrogel within microfluidic chips, as reported previously[1]. Prior to vascularisation, the Polydimethylsiloxane (PDMS) chip was fabricated with a SU-8 Si wafer with 120-μm-thick microchannel patterns as a master mould. A mixture of the Polydimethylsiloxane (PDMS, Dow Corning) and the curing agent in a 10:1 weight ratio was poured to the master mould, and the mixture was cured in an oven at 80°C for 1 hour after degassed. Considering the working distance of ODT, the poured PDMS was adjusted to thickness of 1 mm. After making medium reservoirs and gel ports with 2-mm- and 1-mm-diameter biopsy punches (KAI medical) at the thin PDMS, respectively, they were sterilised by sonication in 70% ethanol for 15 minutes. Plasma (Femto Science Inc.) was treated for 1 minute to bond the patterned surface of the thin PDMS to a 1.5-H-thick- cover glass, and then attach additional two blocks with 4-mm-diameter holes to the unbonded surface of the thin PDMS for reservoirs. The resulting chips were used after restoring hydrophobicity of the bonded surfaces in the 80°C oven for 24 to 72 hours (Fig. 1A).

Red fluorescent protein-transfected human umbilical vein endothelial cells (RFP-HUVECs, Angio-Proteomie) were cultured in a 37°C 5% $CO_2$ incubator in EGM-2MV (Lonza) with 1% Antibiotic-Antimycotic (Thermo Fisher Scientific Inc.), and the cells of passage from 4 to 6 were used for the vascularisation. The RFP-HUVECs detached with 0.25% EDTA-trypsin (w/v) were centrifuged at 200 g for 7 minutes to remove supernatant, and the cell suspensions were prepared in thrombin solution (4U / mL) with a density of $12.5 \times 10^6$ cells/mL. The 10 μL cell suspension was mixed with 10 μL fibrinogen (6 mg/mL) at 5 times on ice, and then the mixture was rapidly injected into a gel channel of the chips. The chips containing the cells were then put in the cell culture incubator within a humid chamber for 15 minutes to form a hydrogel. After incubation, two empty lateral channels were filled with EGM-2MV supplemented with VEGF (50 ng/mL, PeproTech). The chips were kept in the cell culture incubator and the media were replenished every 24 hours to accelerate vascularisation. All chips were daily observed with ODT until the maturation of vascular networks on day 4. Subsequently, some chips were fixed with 4% Paraformaldehyde for 15 minutes at room temperature for comparative structural analysis, and the remaining chips were utilised in vascular permeability assays without fixation.

**Structural quantification**

The microvascular architectures of the images were analysed using ImageJ with 2D skeletonise plugin (NIH, USA), as reported previously[2]. Statistical analysis was performed with OriginPro Software (OriginLab). Measurements were compared using the Student's two-tailed t-test. All tests with $p < 0.05$ were considered to be statistically significant.

**Phase-based microvascular permeability assay**

Consider a solute flux $J_s = -\pi r^2 l \partial_t \Delta c(t)$ through a cylindrical endothelium, whose height, radius, and side surface area are $l$, $r$, and $A = 2\pi rl$, respectively. $\Delta c(t)$ is a transendothelial concentration difference of the solute molecules over time $t$. Assuming that there is no interaction between the solute molecules, the microvascular permeability coefficient $P$ can be calculated from the Fick's 1st law of diffusion[3],

$$P = \frac{J_s}{A\Delta c} = -\frac{1}{\Delta c}\frac{d(\Delta c)}{dt} \times \left(\frac{r}{2}\right). \quad (1)$$

Assuming $P$ and $r$ are temporally constant, integration of Eq. (1) over $t$ becomes

$$\Delta c(t) = \Delta c_i \cdot e^{-(t-t_i)/\tau}, \quad (2)$$

where $\Delta c_i = \Delta c(t_i)$, $t_i$ the initial diffusion timepoint, and $\tau = (r/2)/P$. In the well-established fluorescent-intensity based method, Eq. (1) can be rewritten based on the Beer-lambert law, which relates the linearity between the change in optical density, $I$, and the number of difficusing molecules, $N$:

$$P = -\frac{1}{N_i}\frac{dN}{dt}\bigg|_i \times \left(\frac{r}{2}\right) \approx \frac{1}{I_i - I_{bg}}\frac{I_f - I_i}{\Delta t} \times \left(\frac{r}{2}\right), \tag{3}$$

where $I_i$, $I_f$, and $I_{bg}$ are initial, final, and background fluorescence intensity of the gel region, respectively[2, 4]. Similarly, note that optical phase delay $\Delta\phi(t)$ in QPI measurement is linearly proportional to the concentration difference $\Delta c(t)$, i.e., $\Delta c(t) = \Delta n(t)/\alpha = (\lambda/2\pi h\alpha)\Delta\phi(t)$, where $\Delta n(t)$ and $\alpha$ are RI difference over $t$ and a RI increment of solute molecules, $h$ is the effective endothelial thickness, $\lambda$ is the light wavelength[5-7]. Using the linear relation of $\Delta\phi(t)$ and $\Delta c(t)$, Eq. (2) can be rewritten as

$$\Delta\phi_d(t) = (\Delta\phi_d)_i \cdot e^{-(t-t_i)/\tau}. \tag{4}$$

Here, the optical phase delay induced by the diffusing dextran molecules is $\Delta\phi_d(t) = \Delta\phi(t) - \Delta\phi_{bg} = \Delta\phi(t) - \Delta\phi(0)$. To linearise the problem, we utilise the fit function of $\Delta\phi_d(t)$ written as

$$\log\{\Delta\phi_d(t)\} = \log\{(\Delta\phi_d)_i\} - (t - t_i)/\tau \tag{5}$$

In our phase-based microvascular permeability assay, $P$ was thus calculated using estimated $\tau$ in Eq. (5) with the manually quantified $r$ in Fig. 3D. As a result, from the fit function of $\Delta\phi_d(t)$ the estimated value of $P$ was $8.874\times10^{-6}$ cm/s with R-square = 0.963.

It should be noted that we calculated $P$ directly from $\Delta\phi_d(t)$ instead of $\Delta c(t)$, although we provided $\Delta c(t)$ in the inset at the upper right in Fig.4C for the sake of helping intuitive understanding on the linear relation of $\Delta\phi_d(t)$ and $\Delta c(t)$. For calculation of $\Delta c(t)$, we utilised $\alpha = 0.149$ mL/g for the dextran solution found in other literature[8], and manually quantified $h$ = 15.602 μm.

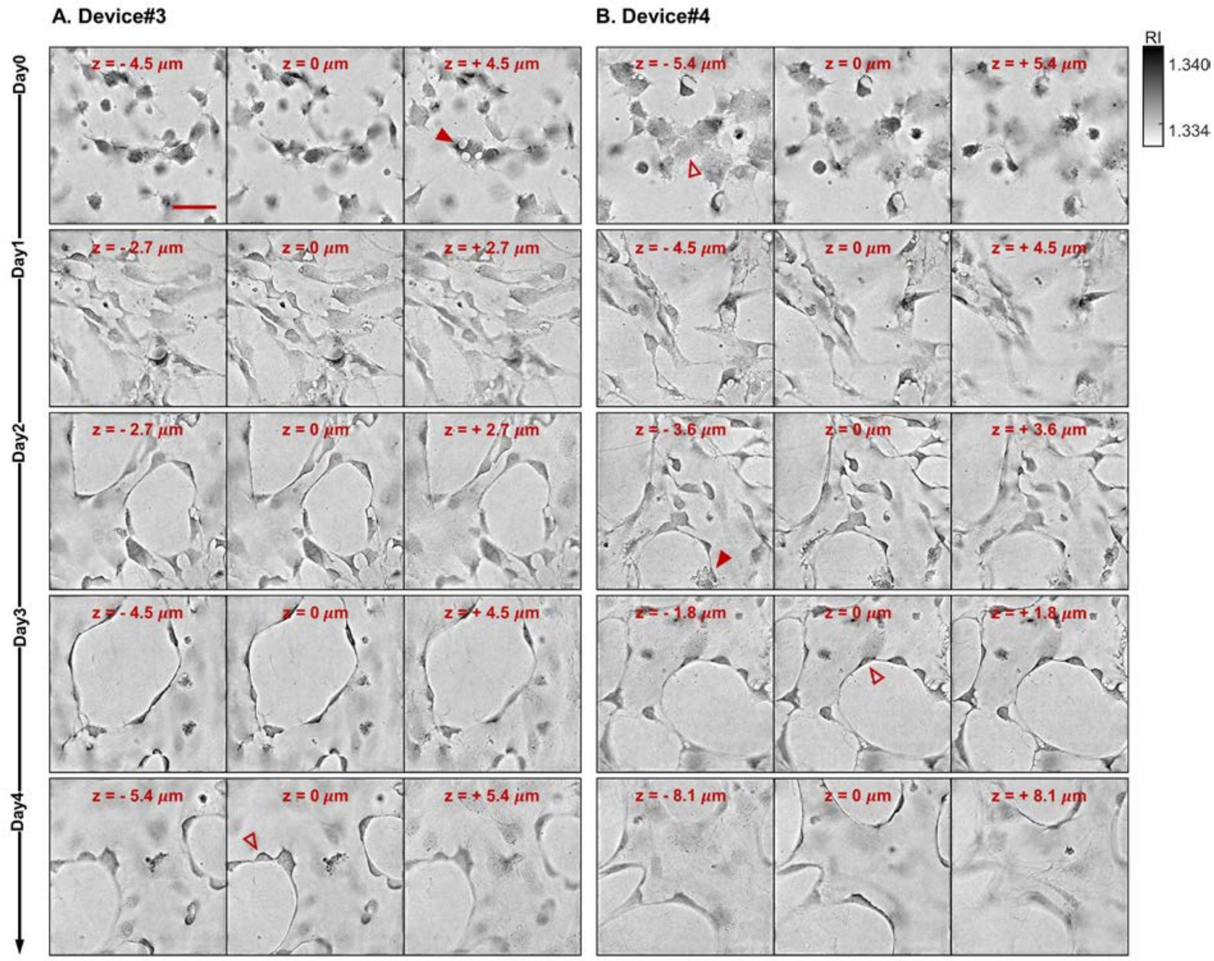


**SI Fig. 1 3D observation of on-chip vasculogenesis using ODT.** (A-B) Multiple XY slice images at 3 z positions selected from 3D RI tomograms of 4-day on-chip vasculogenesis, for two additional devices, A and B, respectively, where various cellular activities (filled arrows) as well as subcellular organelles (open arrows) were observed during vasculogenesis. Scale bar = 50 µm.
